# Human AKTIP interacts with ESCRT proteins and functions at the midbody in cytokinesis

**DOI:** 10.1101/2020.01.19.911891

**Authors:** Chiara Merigliano, Romina Burla, Mattia La Torre, Simona Del Giudice, Hsiang Ling Teo, Chong Wai Liew, Wah Ing Goh, Alexandre Chojnowski, Yolanda Olmos, Irene Chiolo, Jeremy G. Carlton, Domenico Raimondo, Fiammetta Verni, Colin Stewart, Daniela Rhodes, Graham D. Wright, Brian Burke, Isabella Saggio

## Abstract

To complete mitosis, the intercellular bridge that links daughter cells needs to be cleaved. This abscission step is carried out by the sequential recruitment of ESCRT proteins at the midbody. We report here that a new factor, named AKTIP, works in association with ESCRTs. We find that AKTIP binds to the ESCRT I subunit VPS28, and show by high resolution microscopy that AKTIP forms a ring in the dark zone of the intercellular bridge. This ring is positioned in between the circular structures formed by ESCRTs type III. Functionally, we observe that the reduction of AKTIP impinges on the recruitment of the ESCRT III member IST1 at the midbody and causes abscission defects. Taken together, these data indicate that AKTIP is a new factor that contributes to the formation of the ESCRT complex at the midbody and is implicated in the performance of the ESCRT machinery during cytokinetic abscission.

## Introduction

To complete cytokinesis, cells need to cleave the intercellular bridge, a membrane structure enriched in microtubules linking the two daughter cells. This cleavage step, named abscission, is operated by the endosomal sorting complex required for transport (ESCRT) (1, 2). The core of the ESCRT machinery is divided into four subfamilies, ESCRT type I, II and III and VPS4. ESCRT subunits are sequentially positioned at the midbody of the intercellular bridge according to a precise spatiotemporal scheme. The complex of type I ESCRTs composed of the subunits TSG101, VPS37, VPS28 and MVB12 or UBAP1, is recruited first. VPS28 bridges the ESCRT I complex to ESCRT type II, that includes VPS36 and VPS25. In an alternative to this ESCRT I/II pathway, the protein ALIX is first recruited at the midbody (1, 2). Following ESCRT I/II or ALIX localization, ESCRTs type III, such as ESCRTs CHMP2A, CHMP4B and IST1, are positioned. Finally VPS4, an ATPase that controls the supra-molecular reorganization of ESCRT III subunits, works together with the microtubule severing enzyme spastin to finalize abscission (2).

Beyond controlling abscission, the ESCRT machinery operates also in multivesicular body (MVB) biogenesis and at the nuclear envelope to repair membrane discontinuities (2). The plasticity of the machinery is guaranteed by the plurality of ESCRT members and by the factors associated with the ESCRT complex that direct ESCRTs to different intracellular sites. In cytokinesis, the protein CEP55 is an ESCRT accessory factor that forms a disk at the center of the so-called dark zone of the midbody acting as a platform for the recruitment of ESCRTI/II subunits and for the successive organization of the ESCRT machinery (1–4). A double ring structure formed by septins interacts with ESCRT I TSG101 and in this way contributes to demarcate the positioning of the ESCRT complex, therefore working as another ESCRT associated factor (5).

Furthermore, the same ESCRT subunit can act in multiple pathways. For example, the ESCRT III IST1 has been implicated in both abscission and nuclear envelope sealing (6). However, other members of the ESCRT machinery are site-specific. For instance, CHMP7, an atypical ESCRT III, works only at the nuclear envelope (6).

High resolution microscopy has permitted interpretation at the nanometer scale the spatial and temporal organization of ESCRT supra-molecular complexes formed at the midbody (7–10). From such studies, it has been concluded that the ESCRT I TSG101 forms circular structures in the central area of the midbody, while ESCRT II VPS36 is first organized as a single central cortical ring, then extends towards the constriction site with a secondary cone-shaped element (11). The ESCRT III subunits CHMP2A, CHMP4B and IST1 are first organized as double rings at the two sides of the ESCRTI/II complex. In mid/late stage the organization of these ESCRT subunits evolves into spiral structures with progressively smaller diameters at the constriction site (7, 10). The organization of ESCRT III into spirals then leads to the scission of the intercellular bridge by spastin and requires the activity of VPS4 (7, 10).

In our previous work, we characterized a telomeric phenotype associated with a protein named AKTIP in humans (Ft1 in mouse) (12–14). A reduction in AKTIP expression resulted in multiple telomeric signals coupled with the activation of DNA damage markers and with the disorganization of chromatin. In mice, Ft1 reduction causes premature aging defects that are partially rescued by reducing the expression of the DNA damage sensor p53 (15, 16). AKTIP is distributed in the nucleus in a punctate pattern and decorates the nuclear rim in interphase (12, 17). In mitosis, we observed a large AKTIP signal at the midbody (12, 17).

The mechanism by which AKTIP exerts its function is yet to be fully dissected. AKTIP belongs to a subfamily of ubiquitin-conjugating E2 enzyme variants which includes the ESCRT I TSG101 (18). Building on this link with ESCRT I TSG101 and on the fact that AKTIP was detected in at least two ESCRT functional sites (i.e. the nuclear envelope and the midbody), we decided to investigate the function of AKTIP at the midbody from an ESCRT perspective. We performed structured illumination microscopy (SIM) to allow nanometer scale resolution of the localization of AKTIP as was previously done to interpret ESCRT organization and function (7). Here we report that AKTIP forms a ring in the central dark zone of the intercellular bridge in spatial proximity with ESCRT III subunits, and also show that AKTIP binds to the ESCRT I VPS28. We find that AKTIP reduction impinges on ESCRT III IST1 recruitment at the midbody and causes abscission defects, including longer abscission times and binucleation. Taken together, these data provide evidence that AKTIP is a new protein associated with the ESCRT machinery functioning in cytokinesis.

## Results

### AKTIP forms a ring in the dark zone of the intercellular bridge that links the two daughter cells in telophase

To gain insights into the properties of AKTIP we analyzed its spatiotemporal distribution from interphase to late cytokinesis. We immunostained HeLa cells for DNA (DAPI), α-tubulin and endogenous AKTIP. The confocal images show that in interphase, AKTIP is detectable as discrete and abundant puncta at the nuclear rim and within the nucleus (Fig. 1A). During mitosis, the AKTIP signal relocates, and is detected along the microtubules and on the spindle midzone during anaphase. In late telophase, AKTIP accumulates on the intercellular bridge at the center of the midbody (Fig 1A, right panels). At this stage, the AKTIP punctate signal is also visible at the reforming nuclear rim and in the nucleoplasm.

**Figure 1:**
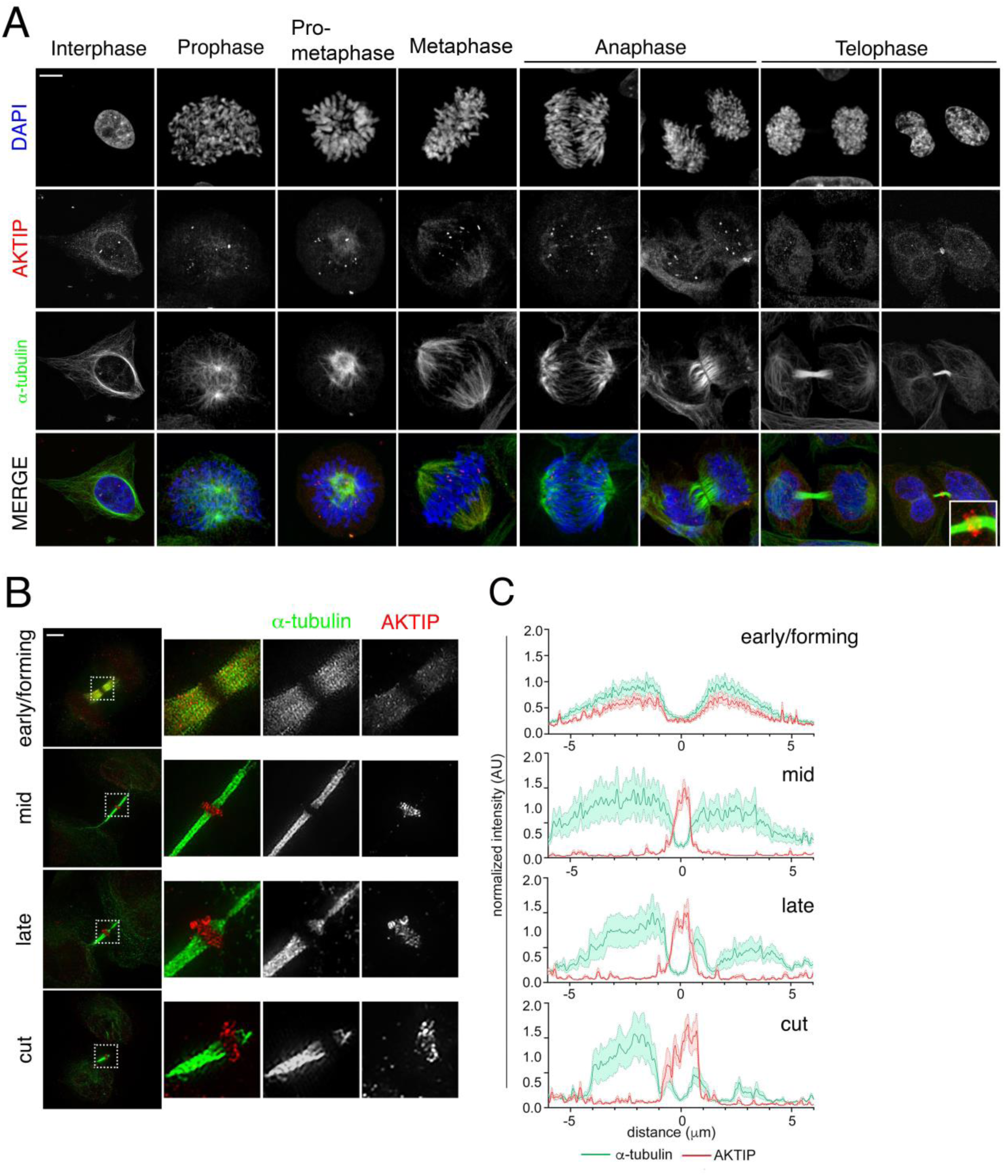
AKTIP localizes at the dark zone of the midbody. **(A)** Confocal immunofluorescence images showing the distribution of endogenous AKTIP in HeLa cells during mitosis and cytokinesis. HeLa cells were stained with anti-AKTIP (red), anti-α-tubulin (green) and DAPI to visualize DNA (blue). Scale bar, 5μm. **(B)** Representative 3D-SIM images of cells with different structural organization observed for AKTIP at the midbody in early, mid, late and cut stages. HeLa cells were stained with α-tubulin (green) and AKTIP (red) antibodies and imaged using 3D-SIM. Panels show the 3D reconstruction including a zoomed-in. Scale bars, 2μm. **(C)** Quantification of signal distribution in representative midbody at different stages.

To validate the specificity of the localization of AKTIP to the midbody, we compared the immunostaining using the monoclonal anti-AKTIP antibody (2A11 WH006400M2 Abnova, Fig. 1 and Fig. S1A upper panel) to that obtained with a polyclonal antibody (HPA041794 Sigma; Fig S1A lower panel). We also analyzed the localization of exogenously expressed AKTIP using anti-FLAG antibody in cells transfected with an AKTIP-FLAG expressing construct (Fig. S1B). In all cases, we observe a clear AKTIP signal at the center of the intercellular bridge (Fig. S1A-B). To conclusively prove specificity, we reduced the expression of the endogenous AKTIP by lentiviral mediated RNA interference (Fig. S1C-D) and observe a drop in the AKTIP signal at the midbody (Fig. S1C and quantification in S1E).

To visualize the assembly of the AKTIP structure at the midbody, we analyzed its temporal and spatial distribution by 3D structured illumination microscopy (3D-SIM), which delivers ∼120nm resolution (19). Cells were stained with anti-AKTIP and anti-α-tubulin antibodies and 3D-SIM images were reconstructed (Fig. 1B and supplementary videos S1). We subdivided midbody stages into early, mid, late and cut, as previously described (5, 7). In the early/forming stage, the midbodies have the largest diameter and the tube is symmetric with respect to the central dark zone. In the mid phase, the microtubules are packaged into a structure that is still symmetric with respect to the dark zone but has a smaller diameter. Late stage midbodies are recognizable by their asymmetry and for the presence of the constriction site. In the early/forming stage, AKTIP is detected as multiple spots on the microtubules of the midbody (Fig. 1B). Image quantification shows the similar profiles of tubulin and AKTIP signals (Fig. 1B and C top panels). During the midstage, AKTIP forms a supra-molecular structure around the central section of the microtubule tube, while staining on the midbody arms is almost absent (Fig. 1B-C, second panel and supplementary videos S1). At this stage, the AKTIP signal reaches its maximum intensity in the dark zone of the midbody where tubulin detection is at its lowest. In late midbodies, the AKTIP structure loses its regularity and part of the AKTIP signal relocalizes and is observed at various distances from the bridge area as demarcated by the α-tubulin signal (Fig. 1B, C third panel and supplementary videos S1).

In order to assess the size of the AKTIP structure we measured its diameter in mid, late and cut midbodies (Fig. 2A). The average internal diameter of AKTIP ring is of 1.05±0.03μm, and the external diameter of 1.89±0.077μm (Fig. 2B). When these data are compared to available measurements for ESCRTs and ESCRT associated factors, we notice that the AKTIP ring is similar to that formed by ESCRT I TSG101 and by ESCRT II VPS36, and slightly larger than that calculated for members of the ESCRT III complex (Fig. 2C).

**Figure 2:**
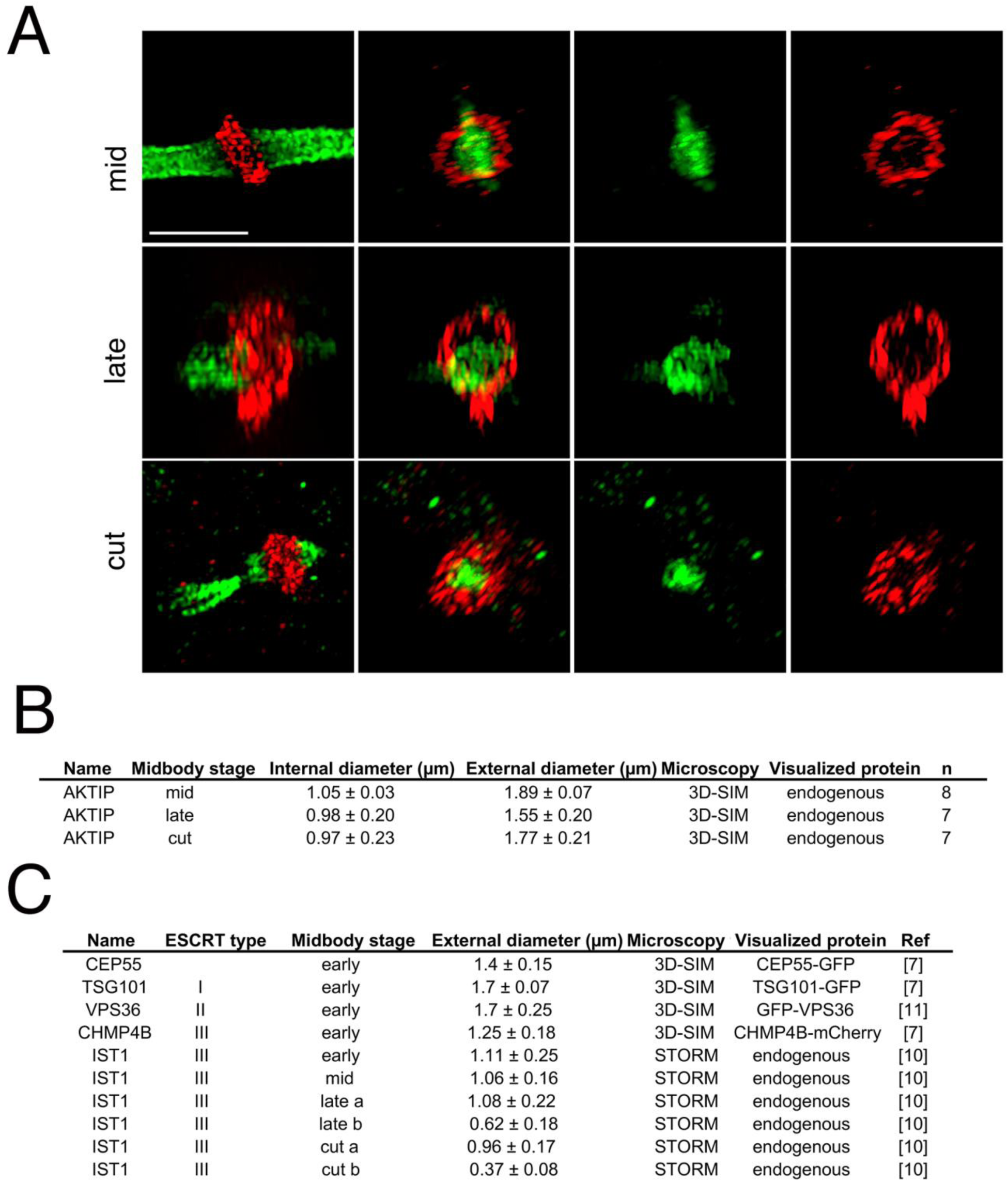
The AKTIP supra-molecular structure is circular and has a diameter similar to that of rings formed by ESCRT subunits. **(A-B)** Representative 3D-SIM images of AKTIP ring and relative measurements (AKTIP showed in red, α-tubulin showed in green). Size of AKTIP structure was measured in mid (n=8), late (n=7) and cut (n=7) midbodies stages. Scale bar, 3µm. **(C)** Size of the diameters of rings formed by ESCRT subunits and ESCRT associated factors at the midbody.

Altogether these results provide evidence that AKTIP forms a supra-molecular ring structure associated with microtubules. The fully formed supra-molecular structure localizes in the dark zone in mid-phase midbodies and is progressively disaggregated in late to cut stage midbodies.

### The rings formed by components of the ESCRT III complex IST1, CHMP4B and CHMP2A flank the central ring formed by AKTIP at the midbody

We then asked whether the AKTIP localization was temporally and spatially linked to subunits of the ESCRT machinery. We focused on the elements of the ESCRT III complex IST1, CHMP4B and CHMP2A, for which the spatial and temporal localization at the midbody had been previously defined (7, 10, 20). In early forming midbodies, a low AKTIP signal follows the tubulin profile (Fig. 3A-B, top panel), and neither AKTIP or IST1 are as yet organized in a supra-molecular structure. In the mid-stage, both AKTIP and IST1 assemble into ring shaped supra-molecular structures. Two IST1 rings flank the central, single, larger and thicker AKTIP ring at the midbody (Fig. 3A-B, second panel and supplementary video S3). In late-to-cut stages, IST1 spirals become apparent on the asymmetric tubulin bridge, and AKTIP progressively loses its circular organization (Fig. 3A-B, bottom two panels and supplementary video S3). AKTIP is absent from the secondary ingression, suggesting it is an early component of the abscission machinery.

**Figure 3:**
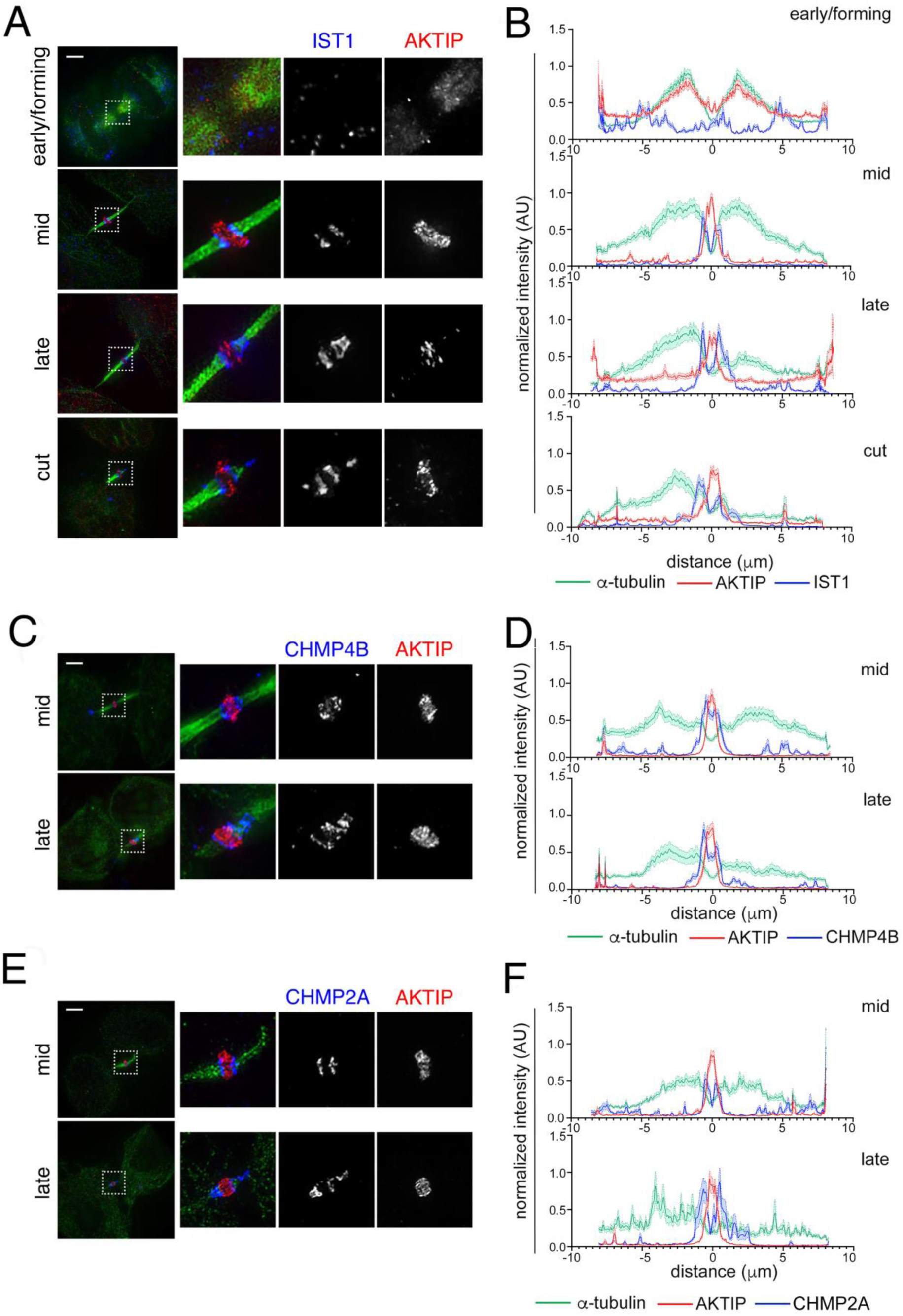
The AKTIP supra-molecular structure is flanked by ESCRT III rings. 3D-SIM images of IST1 **(A-B)**, CHMP4B **(C-D)**, CHMP2A **(E-F)** and AKTIP in HeLa cells. Staining with antibodies against ESCRT III (blue), AKTIP (red) and α-tubulin (green). Scale bar, 2µm. **(B, D, F)** Quantification of signal distribution in midbodies at different stages. (B) early/forming, n=6; mid, n=7; late, n=8; cut, n=6; (D) mid, n=6; late, n=4; (F) mid, n=6; late, n=3.

Subsequently, we analyzed the localization of the two other ESCRT III subunits, CHMP4B (Fig. 3C-D) and CHMP2A (Fig. 3E-F). Co-staining of CHPM2A or CHMP4B and AKTIP shows that the AKTIP ring is sandwiched in between the two rings composed by these ESCRT III subunits in mid-stage (Fig. 3C-D, top panels). In late stages, the ESCRT III subunits CHMP4B and CHMP2A spiral towards the constriction site and AKTIP starts to lose its structural regularity (Fig. 3C-F).

These data show that AKTIP is in the form of a large, circular supra-molecular structure in proximity with the ESCRT III subunits IST1, CHMP2A and CHMP4B, when these get organized into double rings in the central portion of mid-stage midbodies.

### Reduction of AKTIP impairs the recruitment of the ESCRT III member IST1 at the midbody

Taken together our data indicate that AKTIP forms a circular supra-molecular structure in the dark zone, at the center of the intercellular bridge that links the two daughter cells. This AKTIP structure is flanked by ESCRT III subunits in the mid-stage midbody. To provide an understanding on the role played by this AKTIP structure in the context of the ESCRT machinery, we investigated whether AKTIP was required for ESCRT complex assembly. The expression of AKTIP was reduced by RNA interference (Fig. S2) and the presence of ESCRT III subunits at the midbody monitored, focusing on the recruitment of CHMP4B, CHMP2A and IST1. We observe that in cells with reduced AKTIP expression (Fig. S2A), the signal of CHMP4B and CHMP2A at the midbody is only modestly affected (Fig. 4 A, B and D). In contrast to this, the reduction in AKTIP expression impinges significantly on the localization signal of IST1 at the midbody (Fig. 4C-D).

**Figure 4:**
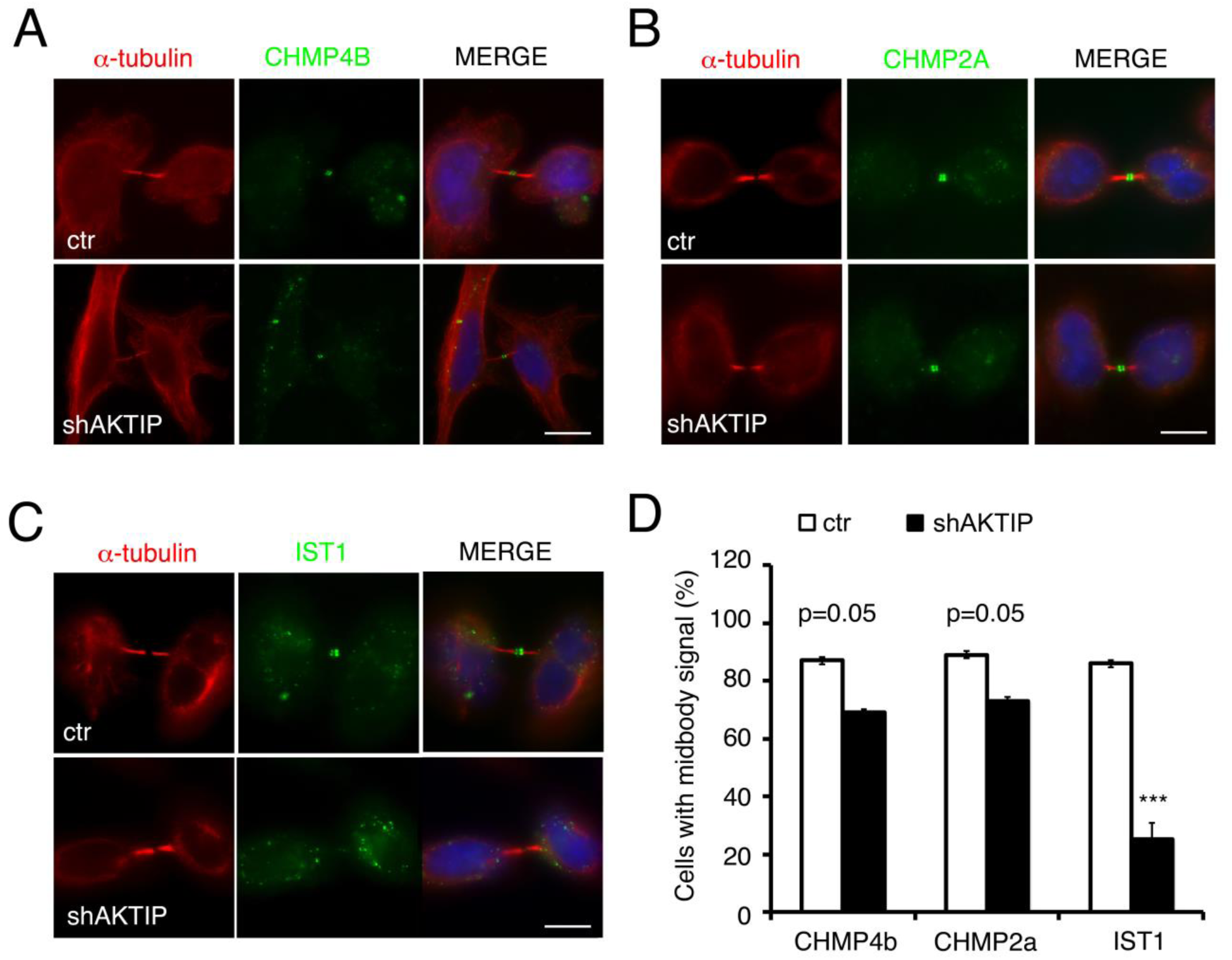
The reduction of AKTIP affects IST1 recruitment to the midbody. **(A-D)** Localization of CHMP4B (A), CHMP2A (B) and IST1 (C) in control and AKTIP reduced (shAKTIP) cells (Figure S2) by immunofluorescence staining with antibodies against CHMP4B/CHMP2A/IST1 (green) and α-tubulin (red). Scale bar, 5μm. **(D)** Quantification of cells from (A-C) showing defects in the localization of IST1 marker at the midbody in cells with reduced AKTIP expression. Results shown are the mean value of three replicates ± SEM. ***p <0.001; Student’s t-test; 100 midbodies per condition were analyzed.

Together these data indicate that, not only AKTIP has a spatial and temporal connection with the ESCRT machinery, but it is also functionally implicated in the correct recruitment of components of the ESCRT machinery at the midbody.

### The reduction of AKTIP expression causes cytokinesis defects including increased abscission time and binucleation

Our results show that the physical proximity of AKTIP with ESCRT III members is paralleled by a role of AKTIP in controlling the localization of the ESCRT III member IST1. IST1 plays a central function in abscission by recruiting the microtubule severing enzyme spastin (21) (22). Spastin is then needed to cut the midbody, coordinating the cytoskeletal and membrane remodeling events necessary to finalize cytokinesis. Given the role exerted by AKTIP in controlling the localization of IST1 at the midbody, we asked whether AKTIP would impact on cytokinesis.

First, we explored whether AKTIP reduction affected cell cycle progression by performing live cell microscopy using HeLa cells stably expressing mCherry tagged α-tubulin. This analysis shows that AKTIP reduction, obtained by RNA interference (Fig. S2B), causes cells to lengthen the abscission stage of cytokinesis from an average time of 107±5min to 169±12min (Fig. 5A-B and Fig. S5 videos). As a consequence of the reduction in AKTIP we observe that cells remain tethered together through their midbody for longer times with respect to control cells (Fig. 5C and supplementary videos S5).

**Figure 5:**
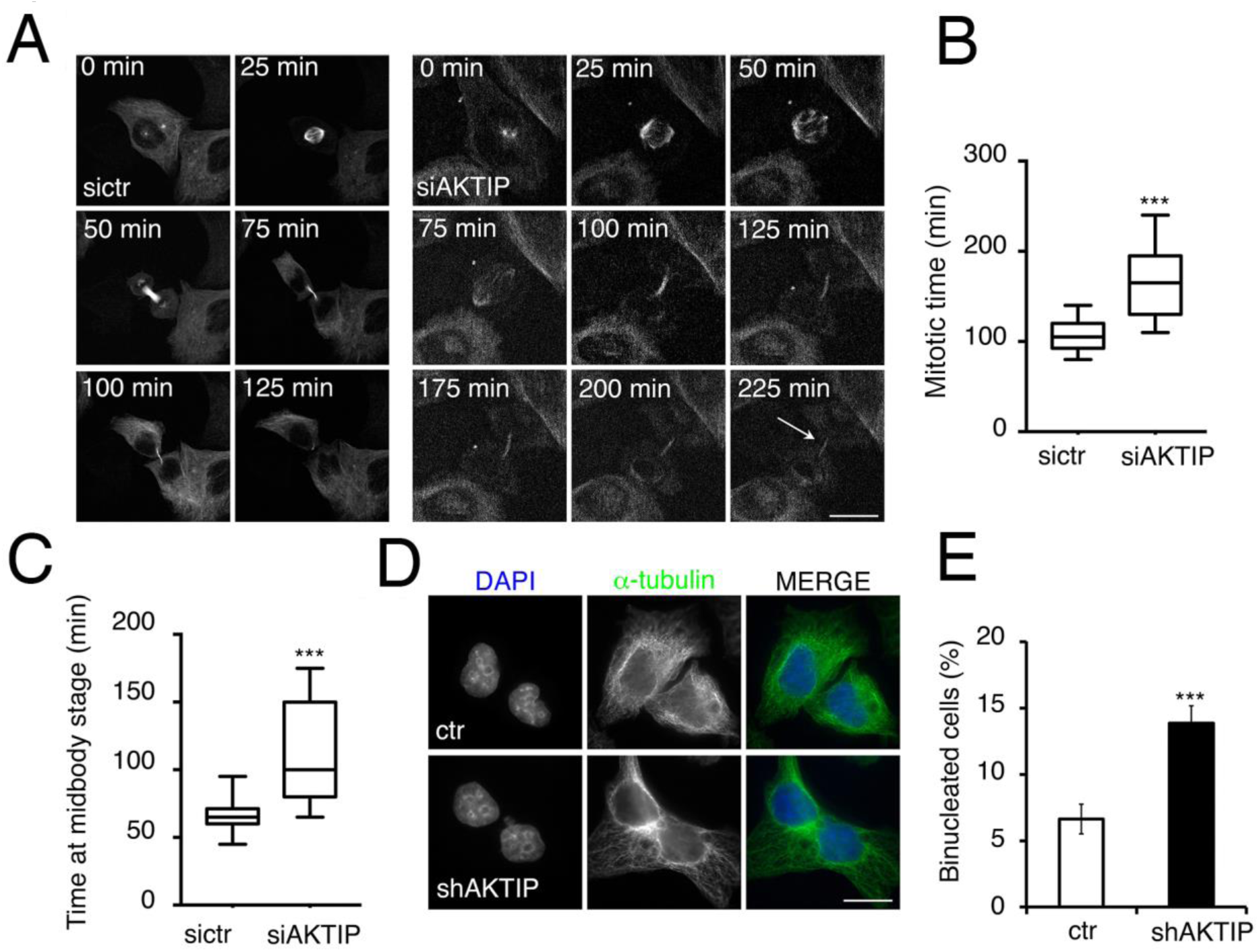
The reduction of AKTIP results in cytokinesis defetcs. **(A)** Selected frames from time-lapse microscopy of HeLa cells stably expressing mCherry-tubulin transfected with ctr (left) or AKTIP (right) siRNA. The arrow points to an example of two cells that remain connected, showing a delay in cytokinesis. Elapsed times are provided in each panel. Movies showing time-lapse images are provided in supplemental materials. Images were recorded every 5min, starting 48hrs after siRNA transfection. **(B-C)** Quantitative analysis of time-lapse microscopy show the time from prometaphase to abscission (B) and from telophase to abscission (C). Results shown are the mean value of two replicates ± SEM. **(D)** Representative images of binucleated cells observed in shAKTIP cells are shown with DAPI in blue and α-tubulin in green. **(E)** Quantification of binucleated cells from (D). Results shown are the mean value of three replicates ± SEM. ***p < 0.001; Student’s t-test. Scale bars, 5μm.

Analyses on fixed cells confirmed that a reduced AKTIP expression causes cytokinesis defects. Specifically, when we immunostained control and AKTIP reduced cells with α-tubulin, compared with control cells, cells with reduced levels of AKTIP show an increase in the number of binucleated cells (Fig. 5D-E and Fig. S2C).

Altogether these data suggest that AKTIP impacts on the IST1 recruitment at the midbody and on the formation of the ESCRT complex, and contributes to the completion of cytokinesis.

### AKTIP interacts with the member of the ESCRT I complex VPS28

Taken together our data demonstrate a physical contiguity of AKTIP with members of the ESCRT machinery at the midbody, an impact of AKTIP on the assembly of the ESCRT machinery, and a role for AKTIP in the completion of cytokinesis. To further understand the possible role of AKTIP as an ESCRT member we searched the 3D structure protein data bank for AKTIP homologues. This analysis identifies TSG101 as an AKTIP homologue with high probability (E-value: 6.3E-7). Significantly, TSG101 is part of the ESCRT I complex, together with VPS37, VPS28 and MVB12 or UBAP1 (1, 2, 23) (24). TSG101 interacts with the ESCRT I member VPS28, which, in turn, bridges the ESCRT I complex to the ESCRT II, and therefore to the ESCRT III complexes. Computer-modeling of AKTIP shows that it superimposes with the UEV domain structure of TSG101 with a root mean square deviation of 1.9 Å (Fig. 6A). Given the structural homology between AKTIP and the ESCRT I TSG101, we asked whether AKTIP was biochemically associated with the ESCRT I complex as is TSG101. To investigate this possibility we carried out a yeast two hybrid screen. Yeast cells were transformed with a plasmid encoding AKTIP fused to the Gal4 DNA-binding domain, in combination with ESCRT I, II, III subunits or associated factors fused to the VP16 activation domain. By measuring LacZ activity in co-transformants we find that AKTIP significantly and selectively interacts with the ESCRT I VPS28 (Fig. 6B). To confirm this interaction in mammalian cells, VPS28 was cloned as a GST-fusion, and AKTIP as HA- or MYC-tagged fusion. 293T cells were co-transfected with control GST or GST-VPS28 and either MYC-AKTIP or HA-AKTIP. Pull down assays followed by Western blotting show that AKTIP interacts with VPS28 (Fig. 6C-D). The interaction is observed with both MYC-tagged AKTIP (Fig. 6C) and with HA-AKTIP (Fig. 6D). GST alone, as expected, does not interact with AKTIP.

**Figure 6:**
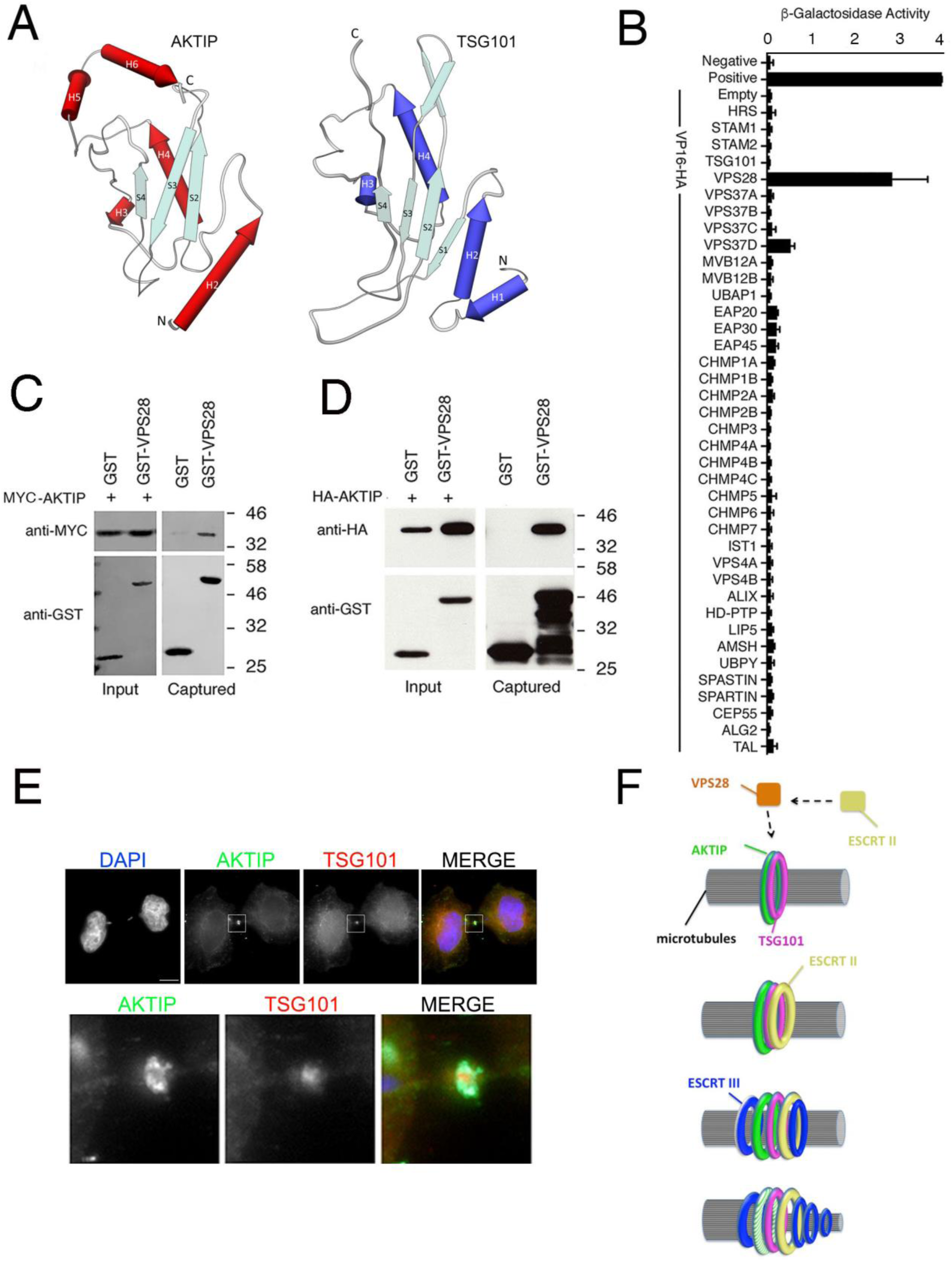
AKTIP has structural similarities with the ESCRT I member TSG101 and interacts with ESCRT I VPS28. **(A)** AKTIP protein 3D model superimposed to TSG101 X-ray solved structure. The model highlights similarities in the central region, and two different elements outside of it: i) the AKTIP central ubiquitin E2 variant (UEV) domain presents two C terminal helices (H5 and H6), absent in TSG101; ii) TSG101 contains two N-terminal helices (H1 and H2) while AKTIP only one (H2). **(B)** AKTIP fused to the Gal4 DNA binding domain was tested for interactions with the human components of ESCRT I, II, III, and ESCRT associated proteins fused to the VP16 activation domain by yeast two-hybrid assay. Error bars indicate the SEM from the mean of triplicate measurements. β-Gal, β-galactosidase; O.D., optical density. **(C-D)** Western blotting showing that AKTIP interacts with GST-VPS28 but not with GST alone. 293T cells were transfected with plasmids encoding the indicated fusion proteins. Purified VPS28-GST or GST alone were used to pull-down interacting proteins; cell lysates and glutathione-bound fractions were then analyzed with MYC or HA antisera as indicated. GST-pull downs were repeated three times. **(E)** Representative images of HeLa cells stained for AKTIP (green) and TSG101 (red) showing that their presence at the midbody is not mutually exclusive. Lower panels, enlargement of regions highlighted by the squares in the upper images. **(F)** Representation of AKTIP implication in the abscission process: AKTIP (green) and TSG101 (purple) form circular structures around microtubules in early/mid stage midbodies. Via an interaction with ESCRT I VPS28 (orange), AKTIP can contribute to recruit ESCRT II (yellow), and ESCRT III (blue). In the final stages of abscission, ESCRT III members evolve into spirals towards the constriction site, while AKTIP is progressively disassembled and absent from the secondary ingression site occupied by ESCRT III.

Taken together, these results indicate that AKTIP interacts with VPS28, which is bound also by TSG101. We then asked whether AKTIP and TSG101 were both present in the dark zone or whether the midbody could contain either AKTIP or TSG101. To this end, we immunostained HeLa cells with both anti-AKTIP and anti-TSG101 antibodies. This analysis shows that 99% (n=100) of the midbodies are positive for both AKTIP and TSG101 (Fig. 6E).

Altogether these data suggest that AKTIP works in association with the ESCRT I complex. In addition, the results point to the hypothesis that AKTIP could contribute to the assembly of the ESCRT machinery at the midbody via its interaction with the ESCRT I VPS28, which can act as a bridge to the ESCRT II, which in turn would impinge on ESCRT III assembly and functional abscission (Fig. 6F).

## Discussion

AKTIP belongs to a subfamily of ubiquitin-conjugating E2 enzyme variants which cannot function directly in the ubiquitination pathway since they lack the cysteine residue required for ubiquitin binding (18). The mechanism of action of members of this family is not yet understood. In the case of AKTIP, we made two observations that suggest that its activity could be associated with the ESCRT complex. A first indication comes from the work by Xu and co-authors, who observed that AKTIP (named FTS) functions in vesicle trafficking (25), a process in which the ESCRT machinery plays a role. The second indication comes from a bioinformatics search in the 3D structure data bank, which indicated that AKTIP has homology with a member of the ESCRT machinery, the ESCRT I subunit TSG101. Here we report that AKTIP is indeed associated with the ESCRT machinery and contributes to its function during abscission.

The first experimental evidence linking AKTIP to the ESCRT complex is based on its supra-molecular organization. We show that AKTIP forms a structure around microtubules at the center of the intercellular bridge where the ESCRT complex is recruited and acts to finalize abscission (5, 7). The supra-molecular structure of AKTIP has the shape of a ring, which in mid phase midbody has an average outer diameter of 1.89μm. This circular organization of AKTIP (showed in figures 1 to 3 and in the related videos) is reminiscent of that of ESCRTs and ESCRT associated factors. These factors form a series of disks and rings in the dark zone, the central area of the midbody, serving as a platform for the successive assembly and structural evolution of the ESCRT machinery. The diameter of the AKTIP ring is similar to that of the ESCRT I member TSG101, which also forms circular structures at the center of the dark zone.

The temporal dynamics of AKTIP at the midbody indicates that the AKTIP supra-molecular assembly happens in early abscission. The assembly of the AKTIP ring is preceded by a phase where AKTIP and tubulin have similar localizations. In the mid-stage of abscission, when the ESCRT III elements have formed full circular structures, the AKTIP ring is found at the center and in proximity with the ESCRT III subunits IST1, CHMP2A and CHMP4B. In the late stages of abscission, when the spiral formed by the ESCRT III factors become evident, AKTIP supra-molecular organization showes a loss of regularity, suggesting that the AKTIP circular structure is needed in early abscission, prior to the final constriction and severing stages.

A further piece of experimental evidence linking AKTIP to the ESCRT machinery is the observation that AKTIP interacts with the ESCRT I VPS28 subunit. Since also the ESCRT I TSG101 also binds to VPS28 (23, 26, 27), we asked whether the presence of AKTIP and TSG101 at the midbody was mutually exclusive. However, both AKTIP and TSG101 are detected simultaneously at the center of the intercellular bridge, which suggests that the regions involved in the interactions between AKTIP and TSG101 with VPS28 are different. This interpretation is consistent with the information obtained by bioinformatic modeling. In fact, while VPS28 binds to the conserved C-terminal region of TSG101 (24), AKTIP differs from TSG101 in that region (see figure 6A), and VPS28 binding sites that are used by TSG101 are predicted to be buried in AKTIP by two C-terminal helices.

Since VPS28 bridges the ESCRT I to the ESCRT II complex (27), the interaction of AKTIP with VPS28 suggests a sequential pathway in which AKTIP via VPS28 is connected to ESCRT II and further then to ESCRT III (see figure 6F). This in turn points to a role of AKTIP in the assembly of the ESCRT III complex. The implication of AKTIP in this process was demonstrated by analyzing the localization of ESCRT III members at the midbody upon depletion of AKTIP. AKTIP impacts the recruitment of ESCRT III complex member IST1, but not on that of ESCRT III CHMP2A and CHMP4B. IST1 is an atypical ESCRT III member that is needed both at the nuclear envelope and in cytokinesis. The fact that an AKTIP reduction impinges more significantly on the ESCRT III IST1 recruitment at the midbody, as compared to its impact on ESCRT IIII CHMP4 and CHMP2B, suggests a further element of distinction of IST1 as compared to the other two ESCRT III elements.

Consistently with the observation that AKTIP impacts on the recruitment of ESCRT III members at the midbody, the reduction of AKTIP expression affects cell division. Cells with lowered AKTIP have significantly longer abscission times and are more frequently binucleated as compared to controls.

In summary, we present evidence that the ubiquitin-conjugating E2 enzyme variant family member AKTIP associates with the ESCRT machinery. AKTIP interacts with the ESCRT I VPS28 and localizes at the midbody, where it forms a ring in proximity to ESCRT III subunits. AKTIP affects the recruitment of the pivotal ESCRT factor IST1 and impinges on cell division (see Fig. 6F). Further work will be required to define the rules for the assembly of AKTIP and other ESCRT factors at the midbody during cell division. It will be also interesting to investigate whether the AKTIP pool localizing at the nuclear envelope plays a role in combination with the ESCRT machinery at the nuclear membrane. In fact, beyond its role in abscission, recent work on IST1 has shown that it is recruited to at the nuclear envelope to seal the membrane of the daughter nuclei by CHMP7 (6). It will be interesting to study whether AKTIP subunits localized at the nuclear envelope are associated with the ESCRT machinery operating at this site and if this activity is related to IST1. In this respect, it is also tempting to speculate that the phenotype of telomere and chromatin damage that we observed in AKTIP depleted cells could be due to defects in the nuclear envelope sealing processes, which would be consistent with the observed phenotype of DNA damage and cell cycle arrest observed in cells with reduced levels of CHMP7 (6).

## Materials and Methods

### Cell culture and RNA interference

HeLa (ATCC CCL-2) and HeLa cells expressing mCherry tubulin (28) were grown at 37°C; 5% CO_2_ in DMEM (Life Technologies) supplemented with 10% FBS (Life Technologies) and 50U/ml penicillin and streptomycin (Life Technologies). For transient RNA interference, cells were cultured in 6-well plates and 20µM siRNA oligonucleotides (Sigma, SASI_Hs01_0086240 for AKTIP, and MISSION® siRNA Universal Negative Control_#1_SIC001 for control, ctr) using Lipofectamine 2000 (Life Technologies) following manufacturer’s protocol. Cells were collected or fixed 72hrs post-transfection. For lentivirus (LV) mediated interference, viruses were produced as previously described (29). The LV-shAKTIP (shAKTIP) and LV-scramble (ctr) vectors were described previously (12). The multiplicity of infection (moi) used was 5pg p24/cell. Transduction was performed in complete medium supplemented with 8µg/ml polybrene (Sigma). After viral addition, cells were centrifuged for 30min at 1800rpm at RT, incubated for 3hrs at 37°C and then transferred to fresh complete medium. Seventy-two hrs post-infection, cells transduced with LVs were subjected to selection in complete medium supplemented with 2µg/ml puromycin (Sigma) and kept under these conditions for further analyses.

### Quantification of gene expression

One-week post-transduction, cells were lysed by addition of TRIzol reagent (Invitrogen) and RNA extracted according to the manufacturer’s instructions. After DNase treatment (Invitrogen), RNA was reverse transcribed into cDNA as already described (30). q-PCR reactions were carried out as previously described (29), using the following primers: AKTIP Forward 5’-TCCACGCTTGGTGTTCGAT-3’; AKTIP Reverse 5’-TCACCTGAGGTGGGATCAACT-3’; GAPDH Forward 5’-TGGGCTACACTGAGCACCAG-3’; GAPDH Reverse 5’-GGGTGTCGCTGTTGAAGTCA-3’ and analyzed with the 2^−ΔΔCq^ method as previously described (31). For Western blotting, 72hrs post-transfection with siRNAs, protein extracts were obtained as previously described (12) and quantified by Bradford assay. 100µg protein extracts were loaded onto pre-cast 4–12% gradient acrylamide gels (Novex, Life Technology). After electro-blotting filters were incubated with anti-AKTIP (HPA041794 Sigma) and anti-actin-HRP conjugated (sc-1615, Santa Cruz Biotechnology) antibodies. Filters were then incubated with anti rabbit HRP-conjugated secondary antibody (sc-2357, Santa Cruz Biotechnology). Detection was performed using the enhanced chemiluminescence system (Clarity ECL, Biorad).

### Microscopy

HeLa cells were seeded onto glass coverslips in 6-well plates and fixed with 3.7% formaldehyde in PBS for 10min. Cells were then permeabilized with 0.25% Triton X-100 in PBS for 5min and treated with PBS 1% BSA for 30min, then stained with primary antibodies in PBS 1% BSA for 1hr at RT. In the case of AKTIP-TSG101 co-immunofluorescence, HeLa cells were seeded on slides, fixed as previously described (5), permeabilized in PBS-0.1% Triton X-100 for 2hrs and successively blocked as described above. The following primary antibodies were used: anti-AKTIP (WH0064400M2 clone 2A11 and HPA041794 Sigma), anti-Tubulin [YL1/2] Rat monoclonal (Abcam, ab6160), anti-CHMP4B (Proteintech, 13683-1-AP), anti-IST1 (Proteintech, 51002-1-AP), anti-CHMP2A (Proteintech, 10477-1-AP) and anti-TSG101 (SantaCruz Biotechnology, sc-7964). Alexa488, Alexa568, Alexa647 or FITC conjugated secondary antibodies were applied in PBS for 45min at RT. Nuclei were visualized using DAPI (4,6 diamidino-2-phenylindole) and coverslips were mounted in Vectashield H-1000. Slides were imaged using Zeiss AxioImager Z1 equipped with a Axiocam 506 monochrome camera. Confocal laser scanning microscopy was performed with Corrsight confocal scanning microscope. Greyscale images were pseudocoloured and combined in Adobe Photoshop CC to create merged images. Live-cell video microscopy was carried out on Corrsight confocal scanning microscope. siRNA-transfected HeLa cells stably expressing mCherry-tubulin were cultured in a 37°C microscope chamber with 5% CO_2_ and observed by phase contrast. Images were acquired every 5min. Images were then analyzed with Fiji (National Institutes of Health, Bethesda, MD).

For 3D-SIM imaging, HeLa cells were seeded onto glass coverslips (high performance coverslips #1.5H, BSF Catalogue #0107052) in 6-well plates and fixed with 3.7% formaldehyde in PBS for 10min at RT and then incubated in 50mM NH4Cl/PBS (15min). Primary and secondary antibodies were applied in PBS-BSA 1% for 1hr at RT and washed in PBS. Acquisition was performed using a DeltaVision OMX v4 Blaze microscope (GE Healthcare, Singapore) with the BGR-FR filter drawer for acquisition of 3D-SIM images. Olympus Plan Apochromat 100×/1.4 PSF oil immersion objective lens was used with liquid-cooled Photometrics Evolve EM-CCD cameras for each channel. 15 images per section per channel were acquired with a z-spacing of 0.125μm (32, 33). Structured illumination reconstruction and wavelength alignment was done using the SoftWorX software (GE Healthcare). 3D volume reconstructions and movies generation were done in Imaris (Bitplane). Image analysis and quantification was performed using Image J (34), Excel (Microsoft) and Prism (Graphpad) software. Fluorescence intensity values represent the average fluorescence intensity measured from a 2.7µm wide band along the axis of the tubulin bundle.

### Yeast Two-Hybrid Assays

Yeast two hybrid assays were performed as previously described (28). Briefly, yeast Y190 cells were co-transformed with plasmids encoding the indicated proteins fused to the VP16 activation domain (pHB18) and AKTIP fused to the Gal4 DNA-binding domain (pGBKT7). Co-transformants were selected on SD-Leu-Trp agar for 72hrs at 30°C, harvested, and LacZ activity was measured using a liquid β-galactosidase assay employing chlorophenolred-ß-D-galactopyranoside (Roche) as a substrate.

### GST pull down

GST pull down were performed as previously described (35). VPS28 was cloned as a GST-fusion into pCAGGS/GST. 293T cells were co-trasfected with 1μg per well of 6-well plate of either pCAGGS/GST or pCAGGS/GSTVPS28 and with with 1μg per well of 6-well plate of pCMV6-Entry-AKTIP-Myc-Flag (ORIGENE) or with 1μg per well of 6-well plate of AKTIP-HA (pCR3.1) for 48hrs. Cells were then harvested and lysed in NP40 lysis buffer (150mM NaCl, 50mM Tris pH7.5, 1mM EDTA). Clarified lysates were incubated with glutathione-Sepharose beads (Amersham Biosciences) for 3hrs at 4°C and washed three times with wash buffer (50mM Tris·HCl, pH 7.4, 150mM NaCl, 5mM EDTA, 5% glycerol, 0.1% Triton X-100). Bead-bound proteins were eluted by boiling in 100µl of Laemmli sample buffer, resolved by SDS-PAGE, as previously described (28). Resolved proteins were transferred onto nitrocellulose by Western blotting and were probed with the indicated antibodies in 5% milk. HRP-conjugated secondary antibodies were incubated with ECL Prime enhanced chemiluminescent substrate (GE Healthcare) and visualized by exposure to autoradiography film. The following primary antibodies were used: anti-HA (ABIN100176, Antibodies Online), anti-MYC (sc-789, Santa Cruz Biotechnology), anti-GST (10000-0 AP, Proteintech). The secondary antibodies used were goat anti-rabbit HRP-conjugated (Cell Signaling).

### Statistics

Statistical analyses were performed using Excel and Graphpad Prism software. Results are shown as mean ± SEM or SD. Data were analyzed using unpaired two-tailed Student’s t-test. p-values below 0.05 were considered significant.

## Acknowledgments

This work has been supported by PRF 2016-67, Progetti di Ricerca, Sapienza University of Rome (RP1181642E87148C) to IS, FIRC (22392), CIB and Fondazione Buzzati Traverso to MLT and IS; Avvio alla Ricerca, Sapienza University of Rome (AR2181642B6F2E48, AR1181642EE61111) to RB, SDG and IS. CM has been supported by EMBO ST fellowship 7621, Veronesi TG 2019. IC is supported by R01GM117376 and NSF Career 1751197. JGC is supported by a Wellcome Trust Senior Research Fellowship 206346/Z/17/Z. The A*STAR Microscopy Platform is supported by the NRF-SIS grant (NRF2017_SISFP10), core funding from A*STAR and an IAF-PP grant (SRIS@Novena). This work is in the memory of P. Bianco.

## Authors’ contributions

CM, RB, MLT, SDG, LTS, LCW, WIG, AC, YO, JGC performed the experiments. IC, JGC, DR, FV, DR, GDW, CS and BB and evaluated the data. IS designed the experiments and wrote the paper.

## Supplementary Information

**Figure S1.**
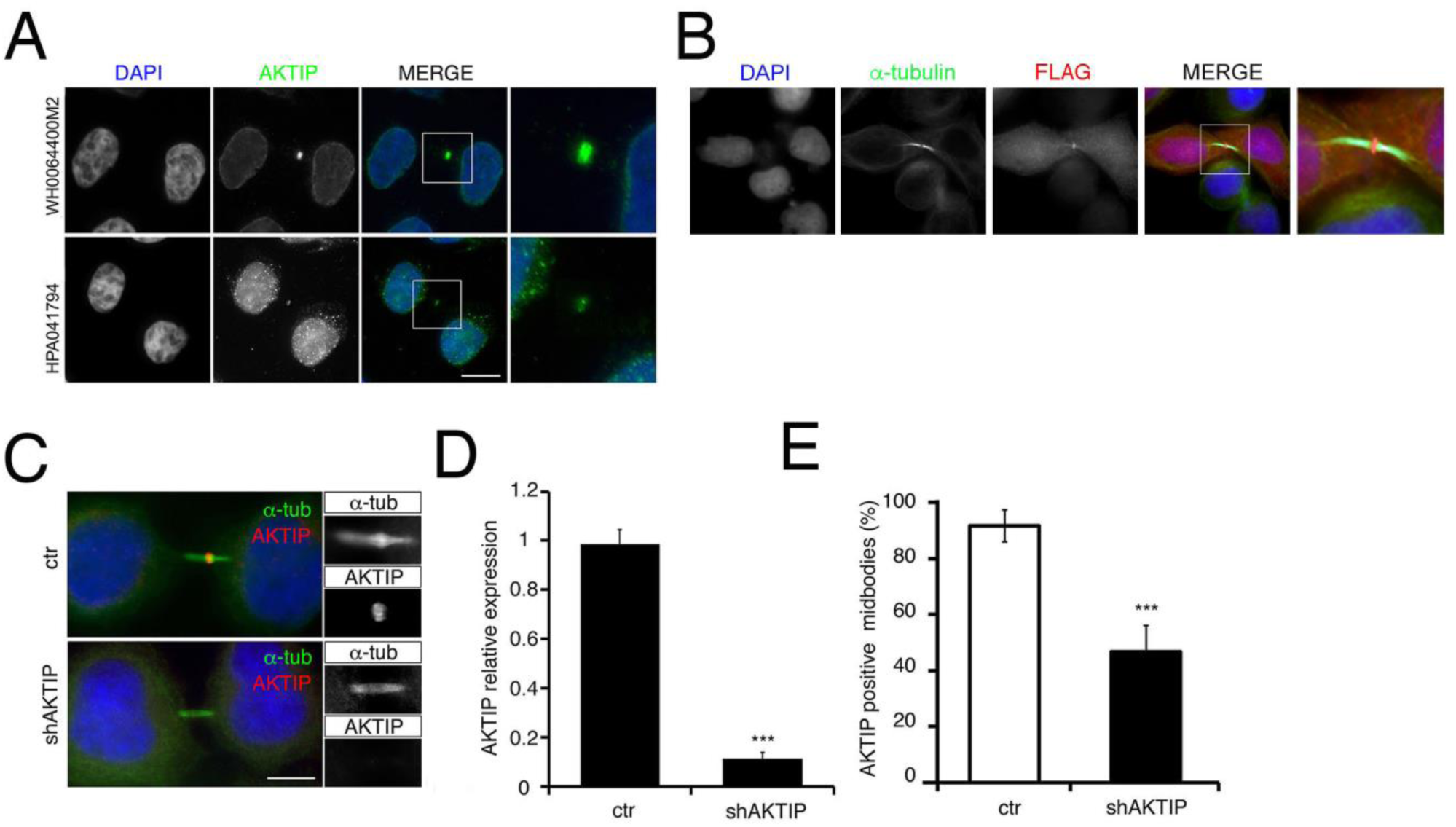
Specificity of the localization of AKTIP at the midbody. **(A)** AKTIP localization at the midbody by immunofluorescence in HeLa cells using anti-AKTIP antibodies WH0064400M2 clone 2A11 (top panel) and HPA041794 (lower panel). **(B)** Detection of exogenous AKTIP-FLAG by immunofluorescence using anti-FLAG antibody. **(C-E)** Immunofluorescence with anti-AKTIP (WH0064400M2 clone 2A11) (C) and qPCR (D) showing that AKTIP reduction causes a drop to 47% AKTIP positively staining midbodies as opposed to 91.7% of control cells (E). Results shown are the mean value of two replicates ± SEM ***p < 0.001; Student’s t-test; 60 midbodies per condition were analyzed. Scale bar 5μm.

**Figure S2.**
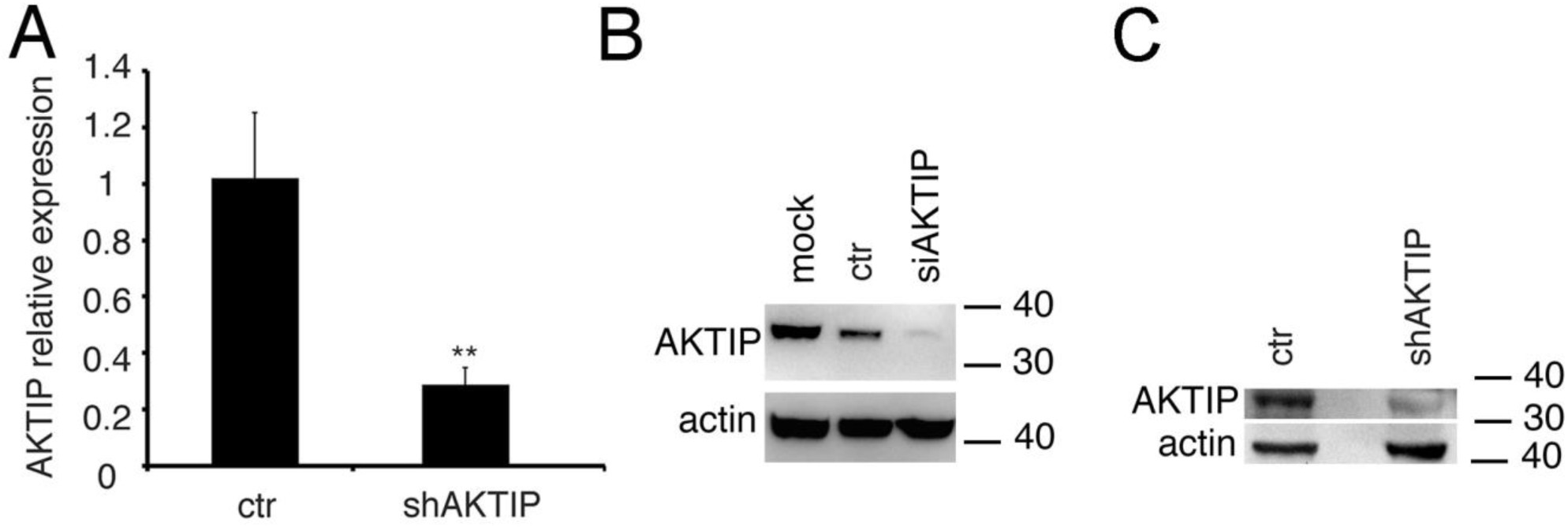
Reduction of AKTIP expression by RNA interference. **(A,C)** qPCR (A) and Western blotting (C) quantification of AKTIP expression in HeLa cells transduced with shRNA directed towards AKTIP with respect to control transduced cells. **(B)** Western blotting showing loss of AKTIP in HeLa cells transfected with siRNAs. Actin was used as loading control.

## Supplementary videos

**S1 videos related to figure 1 (A-B)** 3D volume rendering of mid (A) and late stage (B) midbody imaged with 3D-SIM. HeLa cells were stained with antibodies against α-tubulin (in green) and AKTIP (in red). The volume rendering and the movie generation were performed with IMARIS software (Bitplane).

**S3 videos related to figure 3 (A-B)** 3D volume rendering of midbody imaged with 3D-SIM. HeLa cells were stained with antibodies against α-tubulin (green), IST1 (blue) and AKTIP (red). (A) Mid-stage midbody; (B) late stage midbody. The volume rendering and the movie generation were performed with IMARIS software (Bitplane).

**S5 videos related to figure 5 (A-C)** Time-lapse microscopy of HeLa cells stably expressing mCherry-tubulin treated with siAKTIP (A-B) or ctr siRNA (C). The movie generation was performed with Fiji software.

